# Testable Clinical Signatures for Go-or-Grow Dichotomy of Gliomas along Anisotropic White-matter Tracts

**DOI:** 10.64898/2026.07.01.735477

**Authors:** Sounak Sadhukhan, Debarpita Santra, Souvik Dey

**Affiliations:** Department of Computer Science and Engineering, Techno International New Town, Kolkata, 700156, West Bengal, India; Amity Institute of Information Technology, Amity University Kolkata, Kolkata, 700135, West Bengal, India; Department of Biotherapeutics Research, Manipal Academy of Higher Education, Manipal, 576104, Karnataka, India

**Keywords:** Glioma Invasion, Go-or-Grow, Fisher–KPP, Phenotype Switching, Cell Motility, Mathematical Modeling

## Abstract

On the cellular level, glioma cells seem to be either in a “go” state or in a “grow” state. It is observed that the outer edge of the tumor grows over time. A general rule 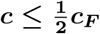 was proven by Thiessen, Conte, Stepien, and Hillen via a substitution argument; however, they weren’t able to derive the exact formula for the tumor’s speed or determine exactly what conditions would cause a tumor to reach its maximum speed. In this article, it is assumed that the cells switch back and forth between moving and growing very fast, and the exact equation for the speed of the tumor expansion is 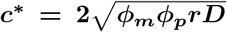. It is found that the same rate of maximum possible growth occurs when cells move for 50% of the time and grow for 50% of the time. If the cells spend more of their time doing one than the other, it grows even more slowly. Moreover, a dimensionless parameter is introduced: 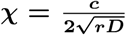, to determine the exact hidden phenotype balance through plugin the speed at which one individual cell moves (***D***), the cell division rate (***r***), and the tumor expansion rate (***c***). The outcome will be near 1 if the old standard tumor model is correct. If it is 0.5 or below, our model is correct; it is either a “go” or “grow” state. In addition, the exact number indicates exactly what percentage of the tumor cells are moving. In the human brain, tumors do not spread evenly in a circle; they spread much faster along with the white-matter tracts apparent through diffusion tensor imaging (DTI). It is shown that our 50% speed limit remains valid along these “brain highways” — a result that has not been previously shown in a mathematical study. Lastly, we address a theoretical question that was left unanswered by past studies using computer simulations and discuss the implications for clinical interpretation of MRI growth rates from real patients using our results.

## 1 Introduction

Gliomas are the most common primary brain tumors, and their high-grade variety (glioblastoma) is almost invariably fatal. The clinical problem is not the bulk tumor; it is the invasion by tumor cells that break away from the main tumor mass and spread diffusely through the brain parenchyma, especially along the aligned fibres of the white matter tracts and the basement membranes of blood vessels. Histopathology consistently shows that these motile cells lie well beyond the MRI-defined border; no matter how aggressive the surgery and focal radiotherapy, they fail to eradicate the invisible invasive rim responsible for recurrence [1, 2]. The speed and shape of the invasion front are therefore central to the disease. Despite significant improvements in cancer treatment, life expectancy remains abysmal, with glioblastoma (GBM) having a median survival of 15 months [3] and only a 9.8% 5-year survival [4]. The poor prognosis is largely due to the extensive spread of tumor cells before diagnosis, and recurrence is almost certain following repeated resections [5]. By contrast, less invasive gliomas have a better prognosis and may be curable [6]. Because of this, the mechanisms of motility and invasion, which underpin GBM resilience through diffuse infiltration, have been investigated. In general, cell migration is initiated by an external *migratory signal*, such as chemical [7], mechanical [8], electrical [9], and other cues. Cells then establish a front where they are moving and a trailing rear. The cell polarises, and actin polymerises at the leading edge, resulting in plasma membrane protrusion. At the cell surface, transmembrane receptors, such as integrins, allow cells to transmit forces generated by interactions between myosin and the cytoskeleton to the extracellular microenvironment. Therefore, manipulation of migratory cues, cytoskeletal filaments, myosin motors, transmembrane receptors, and the extracellular microenvironment are all valuable strategies for inhibiting glioma motility.

In GBMs, cells migrate diffusely through the brain parenchyma, moving along myelinated nerve fibers and the basement membranes of blood vessels (perivascular invasion). GBMs undergo a process known as *Glial-to-Mesenchymal Transition* (GMT) [10]. GBM cells predominantly migrate as individual, solitary cells (mesenchymal or amoeboid single-cell migration). This diffuse, solitary “guerrilla” invasion is exactly the reason why surgeons cannot completely resect GBMs; individual cells drift far away from the main tumor mass into healthy brain tissue. Glioma cells are not alone but rather in a reciprocal relationship with the surrounding microenvironment. They change their surroundings, and the surroundings change them. Glioma progression and invasion are actively dependent on the interaction with the stroma and other non-cancerous cells in the vicinity. Such complex cellular and environmental interactions are not easily captured or explained in traditional biological wet-lab experiments.

In the last 20 years, mathematical models have become an essential and frequently used tool for understanding and simulating these complex tumors. These mathematical models are not purely theoretical; they also have the potential to help in the clinical diagnosis and development of therapeutics. The most popular mathematical model of glioma spread is the Fisher–Kolmogorov–Petrovsky–Piskunov (Fisher–KPP) reaction–diffusion equation [11], consisting of a single tumor-cell density diffusing with coefficient *D* and proliferating logistically at rate *r*. It models the random diffusion and logistic growth of a single population [12]. The Fisher-KPP model is mathematically sound, but its clinical growth rate for tumors is often overestimated. To overcome these drawbacks, mathematicians have been developing coupled systems of partial differential equations (PDEs) and ordinary differential equations (ODEs) that divide cells into a diffusing population and a stationary population [13]. These multiscale models connect subcellular phenomena, including the dynamics of binding of integrins, with macroscopic transport of cells along anisotropic white matter tracts obtained from diffusion tensor imaging (DTI) data [14].

There are unique features and challenges in the mathematical analysis of *Go-or-Grow* (GoG) dichotomy. The travelling wave analysis of the GoG model shows that the maximum invasion speed of a tumor is upper-bounded by the classical Fisher-KPP wave speed, which explains why GoG is a better model for the slow, infiltrative clinical invasion [15]. Mathematicians often reduce the second-order nonlinear reaction-diffusion equations to the first-order Abel differential equations using the Chiellini integrability condition to analytically compute the exact travelling wave forms and tumor growth velocities [16]. A major mathematical challenge for the GoG formulations is diffusion-driven Turing instabilities – also known as a monster on a leash. The resting subpopulation exhibits no spatial diffusivity, so high-frequency spatial perturbations can grow unboundedly, producing complex spatial patterns that resemble GBM heterogeneity but generate severe numerical errors. Moreover, the non-diffusive growing population serves as a biological anchor that decreases the critical domain size required for tumor survival, in contrast to the standard Fisher-KPP models [12]. Time-discretization schemes, such as Rothe’s method, are essential to the proof of the global existence of weak solutions to these complex multiscale PDE-ODE systems, yielding bounded solutions [13].

In the Fisher-KPP reaction-diffusion model, a single tumor-cell density diffuses with coefficient *D* and proliferates logistically at rate *r*. This model is successful clinically, and the speed of its travelling-wave front is the classical Fisher-KPP speed 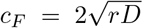 [17]. However, it carries with it a cellular-level, biologically dubious assumption, that the cell is both migrating and proliferating. A large experimental literature, however, backs the GoG dichotomy; *i*.*e*., a cell cannot be proliferating and simultaneously migrating. Giese et al. demonstrated that the motile cells of astrocytoma down-regulate division [18] and vice-versa, and later, *in vivo* imaging studies revealed that single cells within gliomas switch between a motile and a dividing state, with division occurring only during stationary periods [19]. The suggested mechanism is that the cytoskeleton is not able to function in both cytokinesis and locomotion at the same time. This poses a precise modelling question, also in the spirit of this collection, that can be read as a question about cell motility across scales: **what does the GoG trade-off at the microscopic level mean for the macroscopic invasion speed, and is the trade-off sufficiently sharp that it can be distinguished from the single-phenotype proliferation-invasion (PI) model based on measurable quantities?** There have been several studies on GoG reaction–diffusion models and cell-based models; these studies have numerically and asymptotically demonstrated that the spreading speed is modulated by *phenotype-switching* [20–24], and the wave speed has been derived in special cases [21, 23, 24].

From prior mathematical research [25], we already know that, in older mathematical studies, if cells alternate between moving and dividing at a very high rate, we can simplify our equations into a well-known equation (the Fisher-KPP equation). This bedrock was the basis for a recent study by [12] which demonstrated a general rule: tumor growth occurs at most at half the rate predicted by standard models 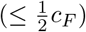, because cells must either move or divide, but not both. To determine that “half-speed”, Thiessen and his colleagues thought it would be easier to put a handy figure into the equations to demonstrate that the speed could never exceed 50%. Although they were only able to write an inequality but unable to determine the precise formula for the tumor’s actual speed and explain under what circumstances within the cells a tumor reaches this maximum speed.

We want to solve that missing problem and make this discovery into a real test tool to use in the lab and in the clinic. In this article, our contributions are multifold. It is shown that the maximum speed of the tumor is reached when exactly 50% of the time is spent moving and exactly 50% growing. A relatively simple formula is developed based on three simple measurements: cell speed, cell division speed, and speed of total tumor growth. By plugging these numbers into the formula, it tells them whether the tumor is following the GoG rule or the old standard model. We showed that this 50% speed limit also applies when tumors spread in complex, directional nerve pathways (called white-matter tracts) in the brain (as seen on MRI). All of these biological tools we backed up with a series of rigorous mathematical proofs, so that it behave correctly in all circumstances.

The **Methods** section includes the underlying two-phenotype reaction–diffusion PDEs, establishes the mathematical well-posedness and cooperative comparison principles of the system and describes numerical integration schemes used to generate the travelling-wave profiles. We then analyze the leading-edge dispersion relation of the two-phenotype system in the **Results** section to get the minimal macroscopic invasion speed (*c*^*\**^), and show that this *c*^*\**^ value is stable in the presence of oscillatory perturbations. A rapid-switching asymptotic reduction is used to establish our central analytical result: if invasive cells switch between the two phenotypes, the GoG dichotomy imposes an exact, closed-form speed ceiling that is exactly half of the classical Fisher–KPP speed, 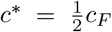; the switching between the two phenotypes is equally divided iff this speed limit is achieved. To verify this speed law and to show how the segregation of phenotypes naturally guides motile cells to lead the invasive rim, direct non-linear numerical simulations are presented. We then convert these theoretical limits to a parameter-less clinical diagnostic ratio (*χ*), which can be used to differentiate between competing invasion hypotheses via macroscopic clinical imaging and further demonstrate that our half-speed scaling envelope remains unchanged for geometry-independent anisotropic diffusion tensor imaging (DTI) white-matter tracts. We integrate the biophysical and clinical implications of our results in the **Discussions**, compare our exact range of attainment with previously published bounds and discuss model limitations. Finally, we conclude this article.

## 2 Methodology

In this research, we have focused on glioma invasion. Here, we consider two subpopulations of tumor cells: *m*(*x, t*) and *p*(*x, t*) denote the densities of motile and proliferative cells, respectively, where *x* denotes the spatial coordinate and *t* denotes time. Let *ρ* = *m* + *p* be the total density of cells. Migratory cells are governed by random motility (Fickian diffusion with coefficient *D*), and proliferative cells obey logistic growth with carrying capacity *K* and growth rate *r*, with cell interconversion rates depending on cell density. The higher the density, the greater the rate of transition to the migratory state; the greater the cell-free space, the greater the rate of transition to the proliferative state. The model is expressed as,

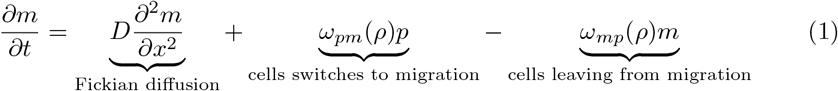

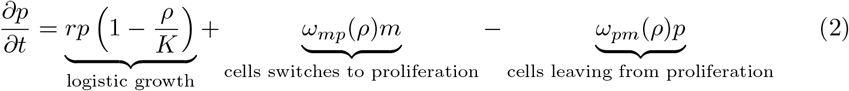

In (1) and (2), *ω*_*pm*_ (*ρ*) denotes the tumour cell switching rate from proliferative to migratory states. In contrast, *ω*_*mp*_ (*ρ*) denotes the switching rate from migratory to the proliferative state. The essential feature of the GoG model is that the diffusion coefficient (*D*) only changes the migratory fraction, and the growth rate (*r*) only changes the proliferative fraction. Fig. 1 illustrates the GoG dichotomy. When a region becomes tightly packed, cells trigger an escape response, shutting down their ability to “go”, and *ω*_*pm*_ (*ρ*) increases with increasing *ρ*. In contrast, a wandering cell hits space (low *ρ*), it senses the available resources, halts migration, and begins to “grow” with increasing *ω*_*mp*_ (*ρ*). The resulting invasion front, a dense proliferative core, gives way to a sparse migratory invasive edge that advances at speed *c*^*\**^ into healthy tissue. We investigate how the two-phenotype-based GoG model (Equations (1) and (2)) behaves on a 1-D spatial domain through two distinct operational frameworks for the switching rates: i) the constant-rate model, and ii) the density-dependent model. In the constant rate model, it is considered that *ω*_*pm*_ (*ρ*) = *α, ω*_*mp*_ (*ρ*) = *β* (*α* and *β* are positive constants). It assumes that cells move between these two teams randomly at fixed linear rates and totally neglects local crowding. It is very amenable to analysis and has analytical information (baselines) that are clean. Case ii) reflects the true biological phenomena using non-linear Hill functions: 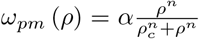 ; switches on sharply once density clears a critical threshold *ρ*_*c*_, and 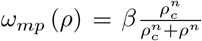 ; shuts-off sharply as density climbs past *ρ*_*c*_. Here, *n* dictates the sensitivity of the switch, *ρ*_*c*_ represents the threshold density where cells become stressed. The escaping behavior due to crowding is modeled by the proliferative-to-migratory rate *ω*_*pm*_ (*ρ*). In contrast, the migratory-to-proliferative rate *ω*_*mp*_ (*ρ*) corresponds to colonizing behavior through free space that is inhibited by a Hill function. The point where these curves intersect is the critical density threshold *ρ*_*c*_ = 0.5, which defines the transition point at which a cell’s dominant phenotype shifts from proliferation to migration. The higher the number of *n*, the closer the GoG decision will be to a binary switch; the lower the number of *n*, the closer the GoG decision will be to a gradual transition. (*For more detail, please refer to the supplementary material section S1*.*1 & S1*.*2*).

**Fig. 1.**
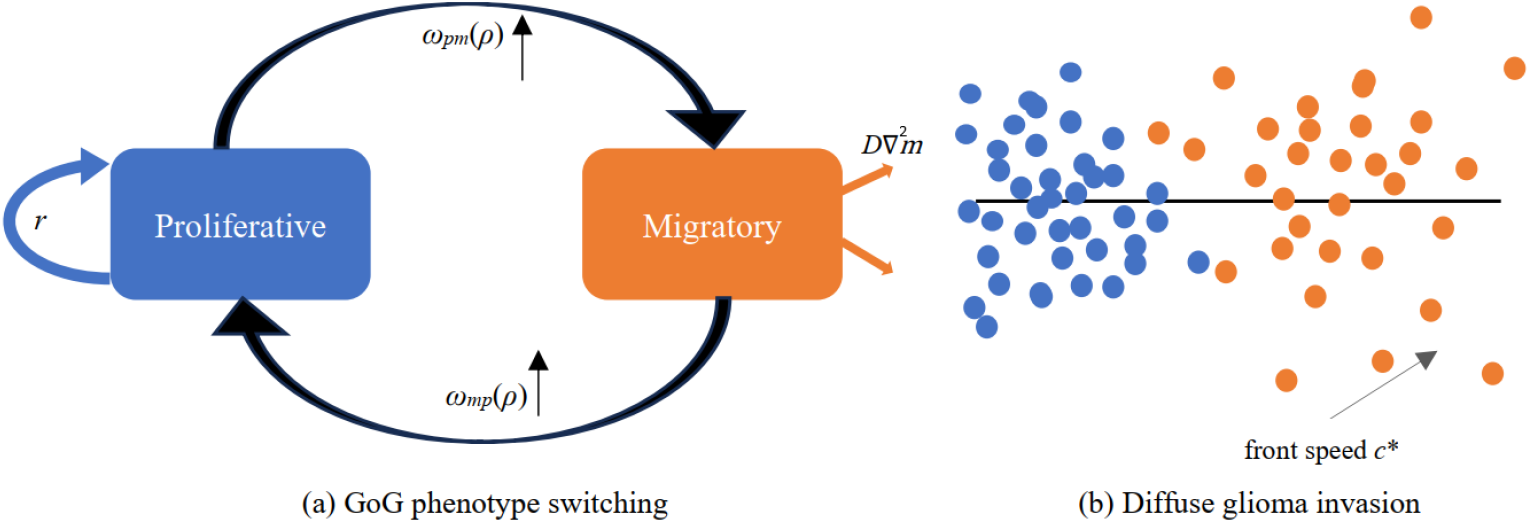
GoG dichotomy and the structure of glioma invasion.

From Equations (1) and (2), we obtained the following 1-D Fisher-KPP Equations (3). This shows that diffusion depends entirely on the moving sub-population and growth depends entirely on the dividing sub-population. The formal limit of the single-phenotype PI or Fisher–KPP model is that all cells move and divide with front speed 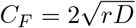 [11, 17].

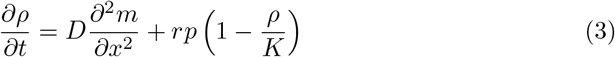

To simplify the system of Equations (1) and (2) and focus on the biological phenomena, non-dimensionalization is applied using the system’s characteristic scales. We define the new dimensionless variables as follows,

1. Dimensionless time: *τ* = *rt*, therefore, 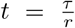
2. Dimensionless space: 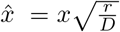, hence, 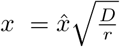
3. Dimensionless densities: 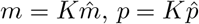, and 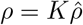

As 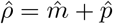. Therefore, Equation (4) becomes,

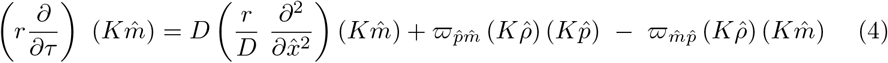

(*For more detail, please refer to the supplementary material section S1*.*3*). Dividing both sides of Equation (4) by the common factor *rK*, we yield the dimensionless Equation (5),

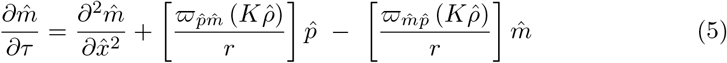

The values or ranges of the model parameters are given in Table 1. By setting the fundamental parameters to unity in Equation (5), and the Fisher Speed (*c*_*F*_): = 2.

**Table 1.** State variables and parameters with representative glioblastoma values.

| Symbol | Meaning | Value/Range | Source |
| --- | --- | --- | --- |
| $m, p$ | migratory, proliferative cell density | $0 - K$ | |
| $D$ | migratory-cell diffusivity | $10^{-3} - 10^{-1} \text{ mm}^2 \text{ day}^{-1}$ (higher in white matter) | [11, 26] |
| $r$ | proliferation rate | $10^{-2} - 10^{-1} \text{ day}^{-1}$ | [11] |
| $K$ | carrying capacity | tissue-dependent | [11] |
| $\alpha$ | proliferative to migratory rate | model parameter | [20, 21] |
| $\beta$ | migratory to proliferative rate | model parameter | [20, 21] |
| $\phi_m = \frac{\alpha}{\alpha + \beta}$ | migratory fraction (fast switching) | $0 - 1$ | derived |

### 2.1 Analysis of the Model

To assess the spatial invasion of the tumor, the mathematical model must be biologically well-behaved and physically realistic. This section examines the behavior of the system in the absence of spatial movement (homogeneous kinetics), establishes the non-negativity and boundedness of the cell populations (the invariant region) and discusses the reason for the movement of the invasion boundary as a “pulled front” controlled by its front.

#### Spatially Homogeneous Steady State

Assuming that the cell populations are spatially homogeneously mixed, no spatial concentration gradients exist. This means that the spatial diffusion term is null 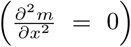. This reduces our system of PDEs to a system of ODEs, which models purely local biological reaction kinetics:

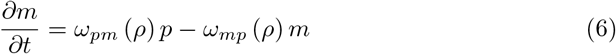

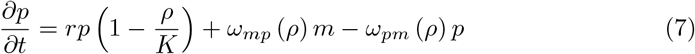

For finding the steady state of this system of Equations (6) and (7), we assign 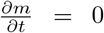, and 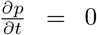. If *m*^*\**^ = 0 and *p*^*\**^ = 0, then both equations are satisfied, corresponding to the tumor-free state.

For a non-trivial population of tumors (population is non-zero), *ρ >* 0 , the process of mass accumulation stops when the logistic growth factor reaches its limit, 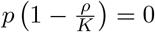. The stable bulk equilibrium state is therefore:

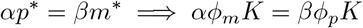

where the fixed phenotype fractions are defined as 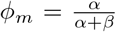 and 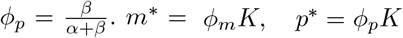 where *ϕ*_*m*_ + *ϕ*_*p*_ = 1

Here, the dynamical system of tumor environment is expressed as a simple linear system as 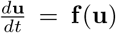, with the cell density vector **u** = (*m, p*)^*T*^ . Linearization is a technique that involves zooming in on a known point of a complex (curved) equation to make the curve appear straight. This approximates the system near the cell densities where it has steady states as a linear system. It forms a matrix of partial derivatives, called the Jacobian matrix, *J*, and determines the stability of two biologically relevant configurations. The first is the evaluation of *J* at the tumor-free state that leads to a characteristic equation *λ*^2^ + (*α* + *β − r*)*λ − βr* = 0 with a strictly positive eigenvalue (*λ*_+_ *>* 0). The positive real part confirms the instability of the healthy state, so that any small amount of inoculum that contains cancer cells will grow exponentially rather than decrease. Secondly, they consider the carrying-capacity state where the total density is at its maximum, *ρ* = *m* + *p* = *K*. Under this crowded equilibrium, the trace (Tr), and the determinant (Det) of the Jacobian 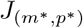 is evaluated and is strictly negative (*i*.*e*., Tr(*J*) *<* 0) and strictly positive (*i*.*e*., Det(*J*) = *rφ*_*p*_(*α* + *β*) *>* 0) respectively, ensuring that all its eigenvalues have negative real parts. This is a confirmation that the density of the tumor core is stable, so that, in time, the cells will naturally reach to a balance and stay at capacity, generating the exact stable mathematical border required to advance a travelling-wave front into healthy tissue. (*For more detail, please refer to the supplementary material section S1*.*4*).

Fig. 2 (a) illustrates the analyzes of a tumor in a closed physical space. A vector field is a way of visualizing the movement of cells as a function of time. It exhibits an unstable tumor-free state at the bottom-left corner (0,0), a very sensitive healthy brain that, as soon as it becomes exposed to a couple of cancer cells, will automatically trigger the green trajectory paths to move upward and in a positive direction. These paths come together safely at a stable bulk equilibrium star without crossing the dashed black line (the invariant region). This represents the mathematical limit at which the number of cells in the tissue can never fall below zero or exceed infinity, due to the tissue’s carrying capacity (*K*). Fig. 2 (b) shows an active movement of the tumor over a tissue domain. It represents a stable invaded front with the total mass (black line) as a wave travelling into space as a rigid body. Importantly, it shows a segregation of the phenotypes at the front - red curve (moving cells), advances of blue curve (dividing cells). This shows that the invading cells are highly motile and then proliferating cells invade behind to form the solid core of the tumor.

**Fig. 2.**
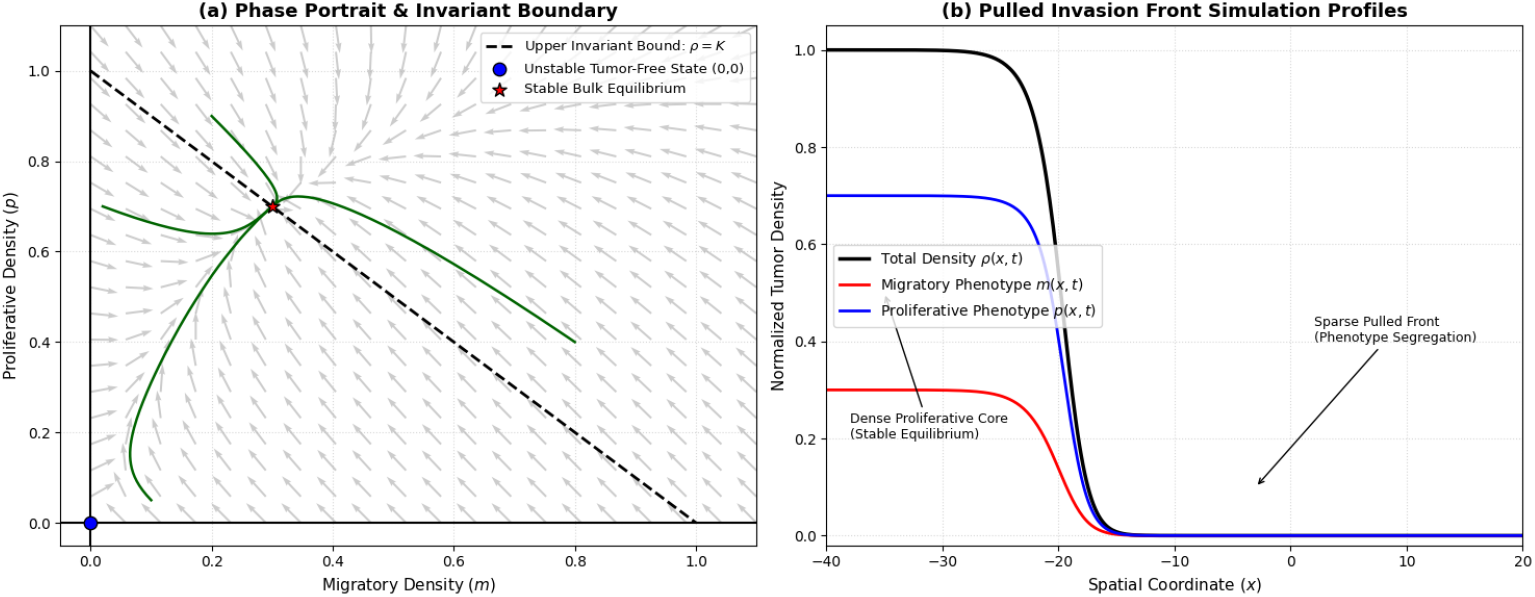
The two-phenotype glioma model is presented in terms of a phase portrait and spatial profiles. The homogeneous reaction kinetics vector field represents the trajectories (green lines) of the system in the mathematically invariant region within the dashed line, which is the carrying capacity line *ρ* = *m* + *p* = *K*. The tumor-free state (0, 0) is an unstable saddle point, and the bulk equilibrium is stable at (*ϕmK, ϕ*_*p*_*K*). The results of (b) are the result of a 1-D numerical simulation of the full partial differential equation system, which shows a stable travelling wave front invading healthy tissue. The models show a clear phenotype segregation at the front of the invasion with motile migratory cells (*m*, red) physically leading the invasion front and proliferating cells (*p*, blue) filling up the invasion behind.

#### Positivity and Comparison

We develop two mathematical characteristics, the positivity and the comparison principle, which are essential to prove the realistic behavior of our spatial system and can be analyzed in a rigorous way to predict invasion speeds at the macroscopic scale. Since it is biologically impossible to have any type of model that predicts a negative number of cancer cells, we do this by showing that our reaction terms are cooperative (or quasi-monotone). The cross-derivatives (off-diagonal terms in the localized Jacobian matrix of the system) are strictly non-negative (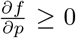 and 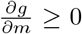), that ensures that the other cell phenotype never slows the growth rate of one cell phenotype. This is a cooperative approach with which we can build an invariant region by using a vector-field boundary argument.

We show that our system is always trapped by the boundary at the physical boundaries (*m* = 0 and *p* = 0 and the ceiling on the total density 0 *≤ ρ* = *m* + *p≤ K*), and that any physically realistic initial data **u**(*x*, 0) *≥* **0** remains bounded in the safe region of 0*≤ ρ* = *m* + *p* ≤ *K* for all future times.

Second, we use this cooperative structure to establish a parabolic comparison principle. It is used in single-variable calculus to determine whether a function is increasing and also in PDEs to guarantee that if one initial profile of tumor is greater than another by ordering the data, then the ordering continues in time as well, **u**_1_(*x*, 0) *≤* **u**_2_(*x*, 0) *⇒* **u**_1_(*x, t*) *≤* **u**_2_(*x, t*) for all time *t*. This principle we explicitly need, as a bounding tool, for “sandwiching” in between a simpler, mathematically tractable linear over- and under-solution of our complex and shifting nonlinear front. We show in this article that the true nonlinear tumor profile cannot outrun or wildly deviate from the absolute leading-edge linearization as *ρ →* 0; this provides a solid mathematical proof for the classification of this as a pulled front. It is precisely this structural verification that allows us the theoretical justification to ignore the intractable nonlinearities of the whole tumor volume and confidently calculate one unambiguous speed of the travelling wave (*c*^*\**^) using only the explicit dispersion relation at the sparse and pioneering edge of the tumor.

## 3. Results

### 3.1 Travelling Waves and the Minimal Invasion Speed

It is considered that the front edge of the tumor is very sparsely occupied. Therefore, the cell density (*ρ*) is close to zero at the far forward position of the tumor. This allows a mathematical simplification: the complex, non-linear “crowding” terms are discarded from the equations, reducing them to a simple, linear set of equations. We assume that this rarefied edge advances as an exponentially decaying wave and show that this assumption leads to a formula called a dispersion relation. The formula is similar: it takes a wave decay rate and outputs the wave speed. The minimum allowed speed on this curve is *c*^*\**^, which is the actual speed of the invading tumor. The original equations (Equations (1) and (2)) are full and very complicated nonlinear equations. The full system makes it almost impossible to obtain a neat, exact algebraic solution for the wave speed. However, it is known that this is a *pulled front, i*.*e*., the front is pulled forward only due to what happens at this zero-density front. Thus, some leading-edge linearization is needed to evade the intractable nonlinear mathematics and to provide an exact, analytical solution for the speed of invasion.

#### Leading-edge Linearization

The *leading-edge linearization* avoids the intractable mathematics in the full nonlinear system and computes the exact speed (*c*^*\**^) of the advancing tumor. These equations are transformed into a moving travelling-wave coordinate system (*ξ* = *x− ct*). Only the absolute forward edge of the tumor where it is very low in density (*ρ→* 0), is considered, and the nonlinear “crowding” terms in these equations are neglected. Then, we tried an exponentially decaying wave guess (*M* (*ξ*) = *M*_0_*e*^*−λξ*^, *P* (*ξ*) = *P*_0_*e*^*−λξ*^) in these simplified equations and arrived at a matrix equation. By assigning the determinant to be equal to zero, we obtain a nice algebraic formula, called a dispersion relation (8):

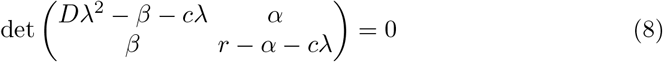

This formula can be used to calculate the wave speed *c* for every spatial decay rate *λ* (Fig. 3). As per the theory of marginal stability, the true macroscopic invasion speed is *c*^*\**^ = min_*λ>*0_ *c*(*λ*), where the minimum is taken over the positive values of *λ*.

**Fig. 3.**
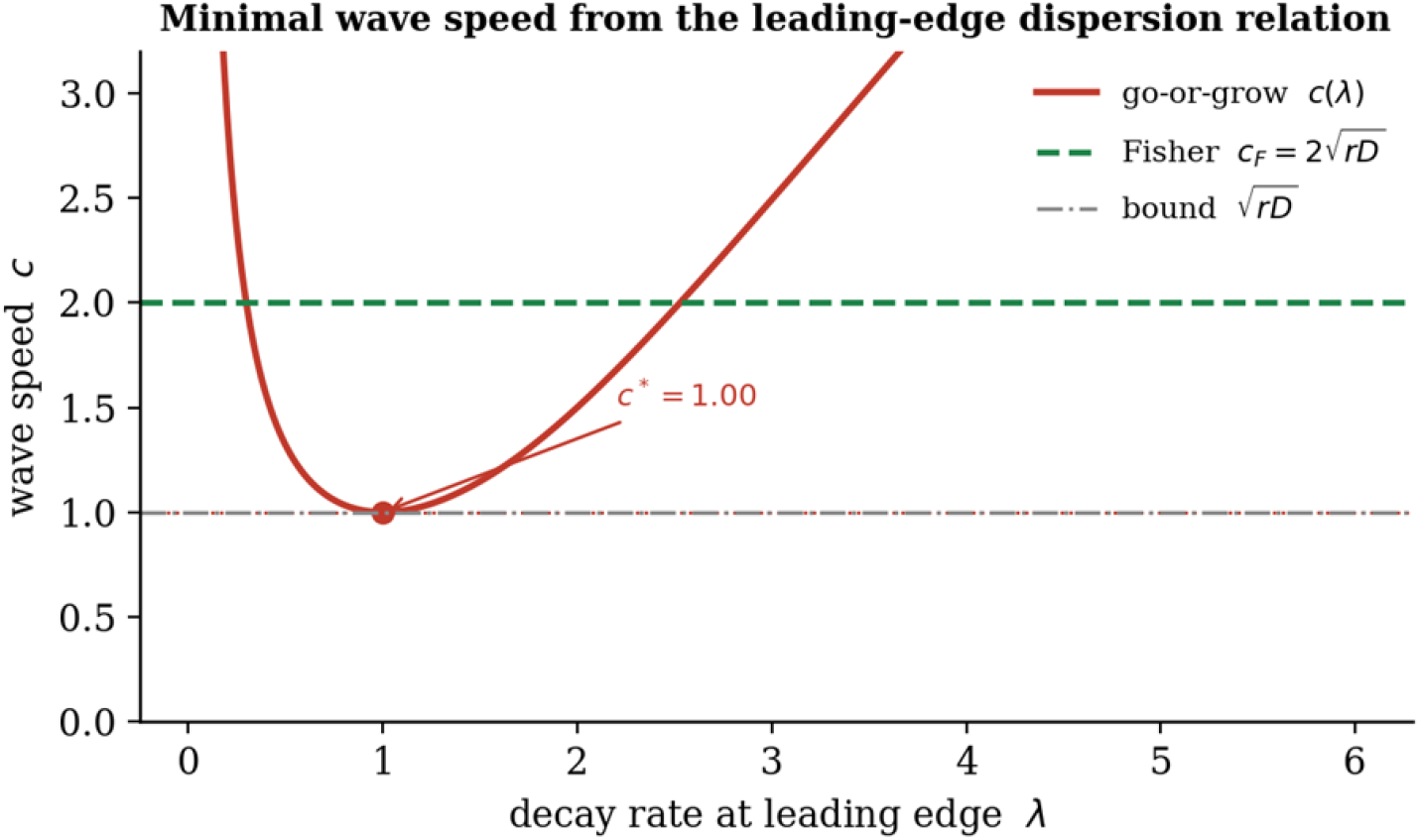
Minimal wave speed from the leading-edge dispersion relation.

#### Simulation of the Invasion Front

We can test the analytical theories using a computational method called the Method of Lines (MOL). This reduces the full, nonlinear PDEs to a coupled system of ODEs that are solved forward in time.

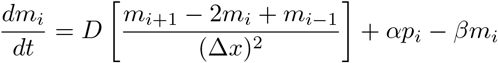

The total cell population moves in the form of a perfectly rigid, stable travelling-wave front, numerically observed speed *c* = 0.94 almost exactly matching their pencil and paper prediction *c*^*\**^ = 0.96 (Fig. 4 (a)); mapping this numerical data into a co-moving coordinate frame (*ξ* = *x* − *c*^*\**^*t*) reveals a fascinating biological phenomenon, called “phenotype segregation”. The leading edge of the advancing wave (as *ξ* → + ∞) has a sudden increase in the local migratory cell fraction (*m/ρ*). At the same time, the core body of the tumor approaches its homogeneous equilibrium value as *ξ → −∞* (*m/ρ* = *φ*_*m*_ = *α/*(*α* + *β*)). This mathematical proof shows that the natural invasion of a tumor, strongly driven by the pioneering action of “go” cells, occurs naturally during the invasion: the “go” cells are highly active and lead the invasion into healthy tissue (Fig. 4 (b)), and the “grow” cells divide behind and are pushed into the dense core of the tumor.

**Fig. 4.**
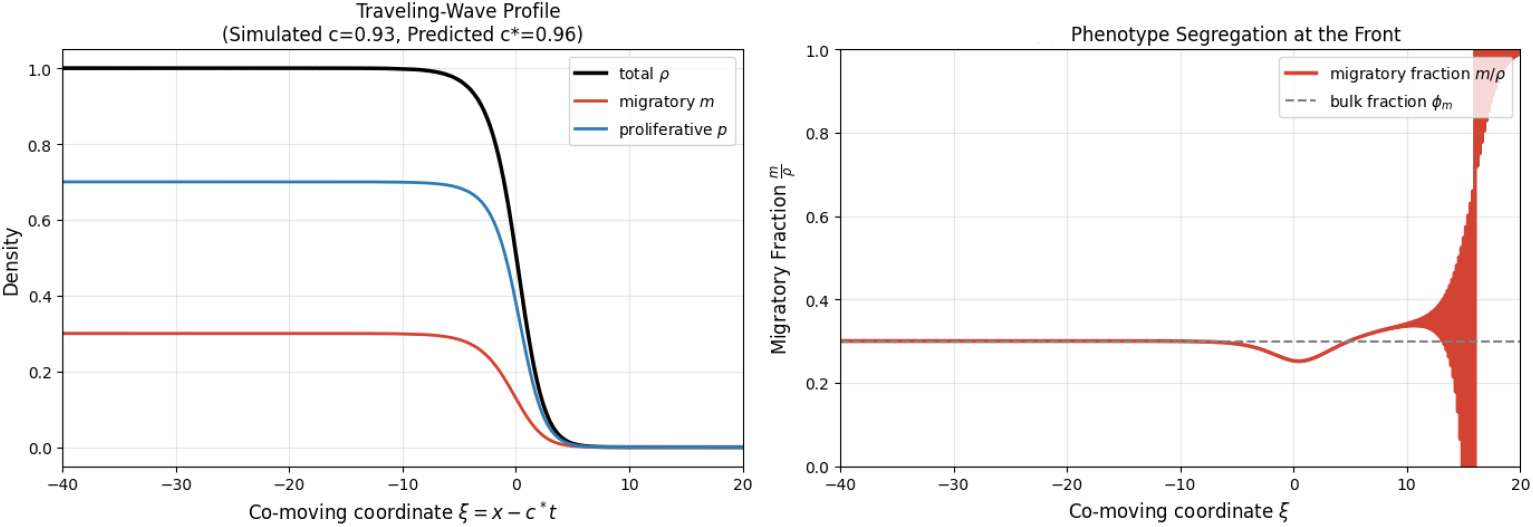
(a) Co-moving profiles of total density *ρ*, migratory *m* and proliferative *p*; the simulated front speed (0.94) matches the predicted *c*^*\**^(0.96). (b) The migratory fraction 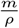 equals the bulk value *ϕ*_*m*_ = 0.3 behind the front but is enriched at the leading edge, so motile cells lead the invasion.

### 3.2 The organising principle: an effective Fisher–KPP limit

In real life, glioma cells switch back and forth between motile and proliferative over the course of just a few hours. But the actual cell division can take much longer (several days). Switching is therefore rapid compared to growth; this separation of timescales is utilized in the classical fast switching reduction of GoG systems [12, 25] to which we appeal below to obtain the sharp testable consequences that are the subject of this paper (*for more detail, please refer to the supplementary material section S2*.*1 & S2*.*2*). Here, we examine the “average” behavior, *i*.*e*., assume that the population of the tumor is a mixture, consisting of a fraction of cells that always behave like motile, *ϕ*_*m*_ and a fraction of cells that always behave like proliferating, *ϕ*_*p*_. To mimic the interplay, a small parameter *ϵ >* 0 is introduced, defined as the ratio of the slow proliferation timescale to the fast switching timescale. The switching rates are scaled with this parameter: 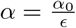 and 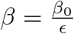. With very rapid switching, these two types of cellular phenotypes are pushed into an “inner balance” proportion. This balance is represented as: *αp* = *βm*, as the total density *ρ* = *m* + *p*. Now, *m* = *ϕ*_*m*_*ρ* and *p* = *ϕ*_*p*_*ρ*, where the phenotypic fractions are defined as: 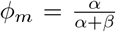 and 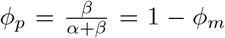. After replacing these fractions (*m* = *ϕ*_*m*_*ρ* and *p* = *ϕ*_*p*_*ρ*) into in Equation (3), we obtain a single closed form Equation (9):

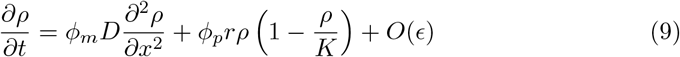

This reveals that in the fast-switching limit with the scaled parameters, the effective diffusivity is *D*_eff_ = *ϕ*_*m*_*D*, and the effective growth rate is *r*_eff_ = *ϕ*_*p*_*r*. Therefore, the spread is due only to the migratory fraction, and growth is due only to the proliferative fraction (10).

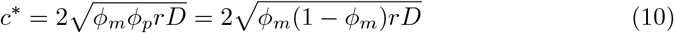

The above claim sharpens the bound of [12], whose proof substitutes a single convenient decay rate into the leading-edge dispersion relation and so establishes 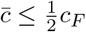 as an inequality without identifying the true minimum or the condition for equality.

Here the minimum is found explicitly by direct calculus on the dispersion relation of Sec. 3.1, giving both the exact minimiser and the attainment condition 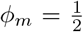: the bound is not merely an upper estimate but is achieved, and achieved only when cells split their time equally between migrating and proliferating.

The cell needs to turn off its motility system when it divides, which means that the interplay between the motile and the proliferative cell must be flawless for the cell to invade as much as possible. The fraction of time a cell is in the “go” state is denoted by *ϕ*_*m*_. The time spent in the “grow” state is (1 *− ϕ*_*m*_). The biological growth rate, or proliferation rate of cells, is represented by *r* and their biological diffusion/motility rate by *D*. Finally, we set the classical “Fisher Speed” 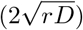 equal to *c*_*F*_.

Let’s define the function *f* (*ϕ*_*m*_) for the variable part of this equation. The critical point, the point at which this function reaches its maximum value, is 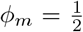. After plugin this optimal fraction 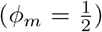 back to the original Equation 10, we get 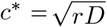. Hence, the fastest a GoG tumor can grow is half the classical Fisher speed, 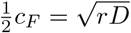, which is the fastest a finite difference solution can grow.

To test the global strength of the speed law of fast switching, we performed a thorough grid sweep of the invasion front over a large range of phenotypic transition rates, *α* and *β*. The simulated front speeds produced show that the highest possible invasion rate is achieved on the diagonal where *α ≈ β* as predicted by theory, thus giving visual proof of the theoretical need for the balanced phenotype distribution to obtain the highest possible invasion rate. Moreover, the absolute error between our theoretical speed equation (*c*^*\**^) and these simulated results shows the wide range of accuracy of the mathematical model. The reduction of the effective Fisher-KPP is very reliable, as the error is negligibly small for almost all of the parameter space in the biological realm. The only points that are significantly different are extreme edge cases, in which a strong migratory bias (high *α* and low *β*) leads to cells spending time in the motile state, a mild deviation from the assumption of the theoretical speed limit, that cells represent a fast switch between two states.

Lastly, what happens when the switching speeds are switched in a realistic situation: cells do not switch back and forth infinitely fast (the “fast switching limit”). If the cells that make up the tumor change states just a little bit slower, then a cell can experience a longer, unbroken period of pure movement or pure division before it must halt. This leads to slightly faster speeds than the strict mathematical limit line indicated in the graphs. Despite all of this, the basic rule remains: The overall invasion speed will never exceed 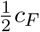 ceiling (Fig. 5).

**Fig. 5.**
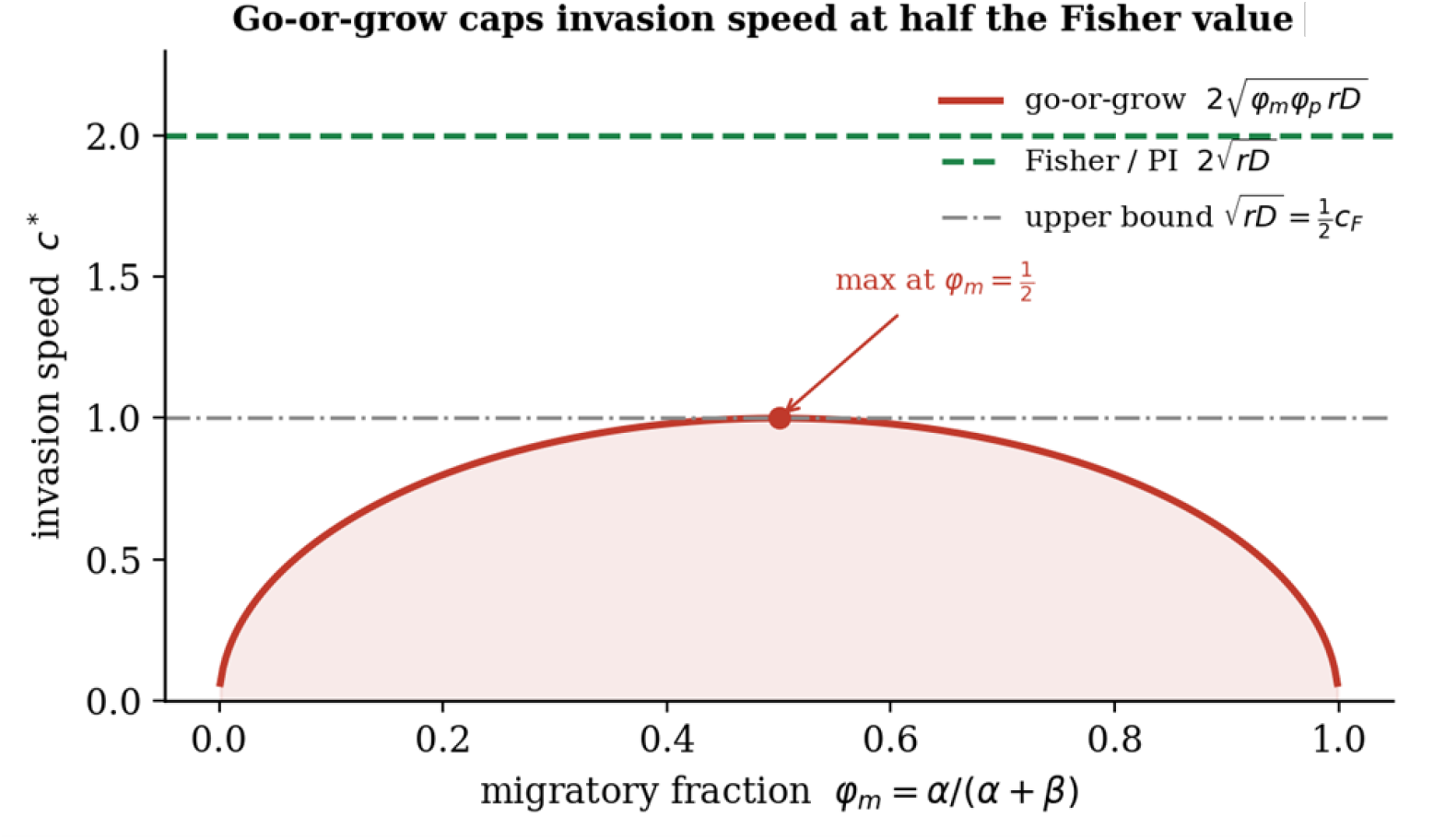
Convergence to the fast-switching law. Exact dispersion-relation speed versus *ϕ*_*m*_ for slow, moderate and fast switching (total rate *T* = *r*, 4*r*, 20*r*) approaches the closed form (Equation 14) (dotted) as switching accelerates. All curves remain at or below 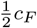 ; slower switching yields slightly faster fronts.

**Fig. 6.**
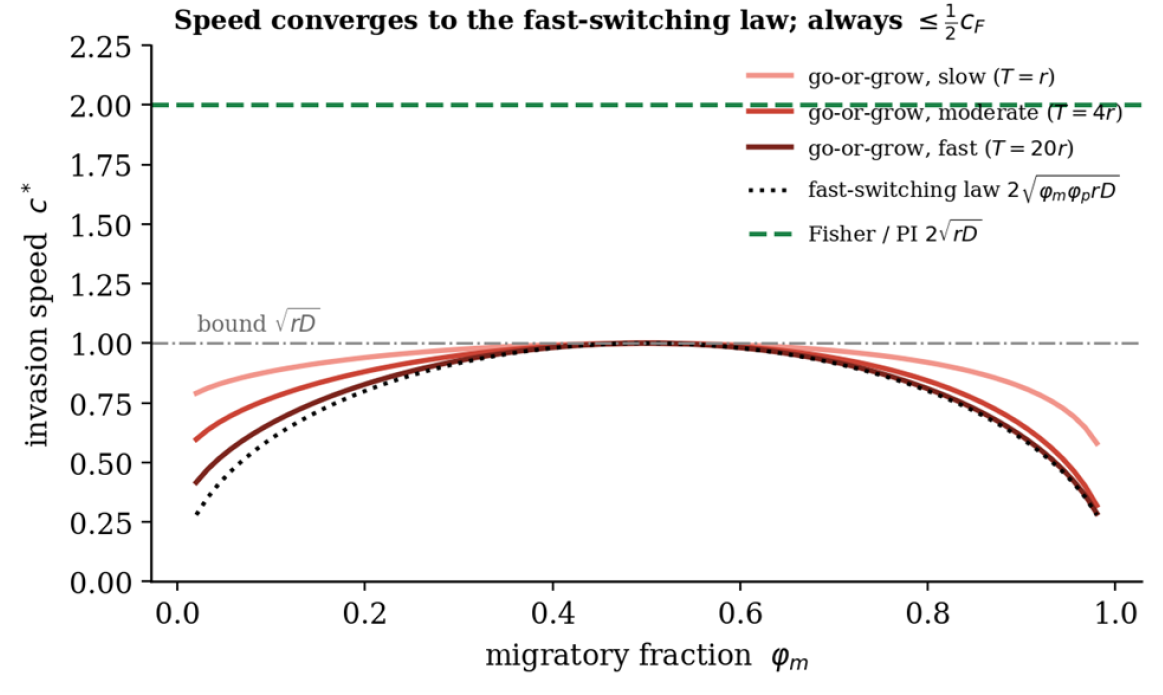
Convergence to the fast-switching law. Exact dispersion-relation speed versus *ϕ*_*m*_ for slow, moderate and fast switching (total rate *T* = *r*, 4*r*, 20*r*) approaches the closed form (Equation (10) (dotted) as switching accelerates. All curves remain at or below 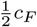; slower switching yields slightly faster fronts.

### 3.3 A Testable Signature that Distinguishes The Hypotheses

The paper describes how the mathematical theories covered above are not only abstract concepts, but also can be directly tested in a laboratory or clinical environment to see how a tumor is, in fact, behaving. Scientists have to be able to measure three things independently:

1. *D* (motility): The speed with which individual tumor cells are moving around, which can be tracked under a microscope in a lab.
2. *r* (proliferation): The speed with which the tumor cells are dividing and multiplying to create new cells.
3. *c* (front Speed): The speed with which the outer edge of the entire tumor is growing and spreading, which doctors can measure by serial brain scans over time.

Researchers set up a dimensionless ratio, *χ* to compare the actual tumor speed with the theoretical maximum. This is given by the measured macroscopic front speed (*c*) divided by the classical Fisher speed limit 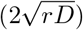:

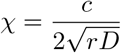

The two biological models give different mathematical predictions for the expected value of *χ*.

PI Model: When cells are migrating and proliferating at the same time, the equation governing the dynamics is the standard Fisher-KPP equation: 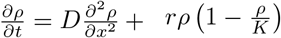. The anticipated speed is 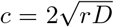. So, according to the PI hypothesis, we can predict:

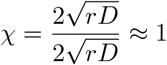

When changing between states is rapid, the system can be reduced to an effective Fisher equation with the speed 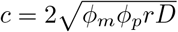. Because the term *ϕ*_*m*_*ϕ*_*p*_ (or *ϕ*_*m*_(1 *− ϕ*_*m*_)) maxes out at 1*/*4, the maximum expected speed is 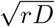. Based on this, the GoG hypothesis predicts that the ratio is kept to a maximum of half:

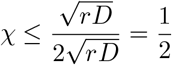

When the ratio measured by the researchers is consistent with the GoG hypothesis 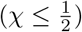, the mathematics enables the researchers to work back to determine the true proportion of migratory cells (*ϕ*_*m*_), without counting them one by one. Substituting the GoG speed equation into the *χ* equation reduces the terms to:

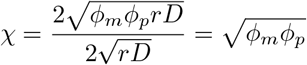

Since the migratory and proliferative fractions must equal to 1 (*ϕ*_*p*_ = 1 *− ϕ*_*m*_), we can rewrite as *ϕ*_*m*_:

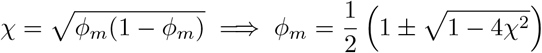

This algebraic derivation demonstrates exactly how a macroscopic speed measurement (*c*), combined with single-cell rates (*D, r*), yields a precise internal calculation of the tumor’s phenotypic balance.

### 3.4 Anisotropic Invasion Along White-Matter Tracts

So far, the spatial domain studied for the GoG dichotomy has been assumed to be homogeneous, and the cell motility isotropic. The microenvironment in the human brain is very complex and heterogeneous, however. Infiltrating glioma cells move in a non-random manner and consistently localize and migrate along the brain’s white-matter tracts, which serve as anatomical “highways”. The clinical and histological observations have consistently been that infiltrating glioma cells do not migrate randomly but rather exploit the brain’s white-matter tracts as highways. This localized guidance results in quick and highly directionally anisotropic invasion that determines the irregular shape of the macroscopic tumor.

To remain relevant to the actual clinical applications, our model needs to take into account the fact that the white matter architecture of the brain is typically quantified by capturing the diffusion of water molecules, which is an indicator of tissue alignment, and is routinely measured in modern neuroimaging, particularly in the form of Diffusion Tensor Imaging (DTI). In this section, we generalize our effective Fisher-KPP limit to a diffusion tensor instead of a scalar diffusion coefficient. In this way, we explore how the tight phenotypic speed limit previously obtained works in a highly directional environment, ultimately showing that the structural cost of mutual exclusivity does not depend on the underlying tissue geometry. (*For more detail, please refer to the supplementary material section S2*.*3*).

- **The Biological Reality (brain “highways”)**: Tumor cells in a simple laboratory petri dish may spread out in an ideal circle, progressing in all directions at the same speed. This is called “isotropic diffusion”. In the human brain, however, glioma cells have anisotropic diffusion; they move at different speeds depending on their direction of motion. The glioma cells are more attracted to the aligned fibers of white-matter tracts than to pushing across them or migrating through the denser grey matter. Therefore, real gliomas are not round, but are star-like or elongated, with projections of the tumor extending along these neurological pathways.
- **Mathematical Upgrade (from scalar to tensor)**: The mathematical model of this needs to be improved from the simple single-number diffusion coefficient (*D*) described in the previous sections. The model uses a diffusion tensor (**D**) in place of the scalar *D*. It states that the motion of the tumor in the north-south direction is fast (along the fibers), while in the east-west direction is slow (across the fibers). If we put this tensor into the migration equation, the new formula for the direction of the speed of the tumor will become Equation (11):

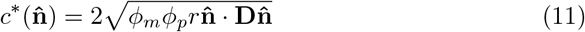

The direction in which we look at it is given by the normal vector 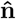, which denotes a specific direction of the spreading front. The vector 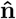 is used mathematically to pull out the specific motility rate for that exact direction from the tensor compass.
- **The core Insight (a universal speed penalty)**: The most important mathematical discovery in this section, *i*.*e*., the GoG penalty is universal and independent of the environment. The penalty factor that we uncovered in the simple model *ϕ*_*m*_*ϕ*_*p*_, is in the anisotropic speed equation. This implies, when the glioma cell is crawling slowly through dense gray matter (slow axis), the maximum speed the tumor front can reach in that direction is half that of what the classical Fisher-KPP model would yield for that same direction; and if the glioma cell is sprinting down a white-matter highway (fast axis), the maximum speed the tumor front can achieve in that direction is still one-half of what the classical Fisher-KPP model would yield in that same direction. The speed envelope for GoG is not “warped” or “distorted” by the geography of the brain, as you can see in your Fig. 7 polar plot - it is just a perfectly scaled-down, miniature copy of the older model’s predictions. Biological limitation is an intrinsic barrier to the cell; it is essentially a physical limitation in the cell that means the tumor is limited, no matter what happens in the outside terrain.

**Fig. 7.**
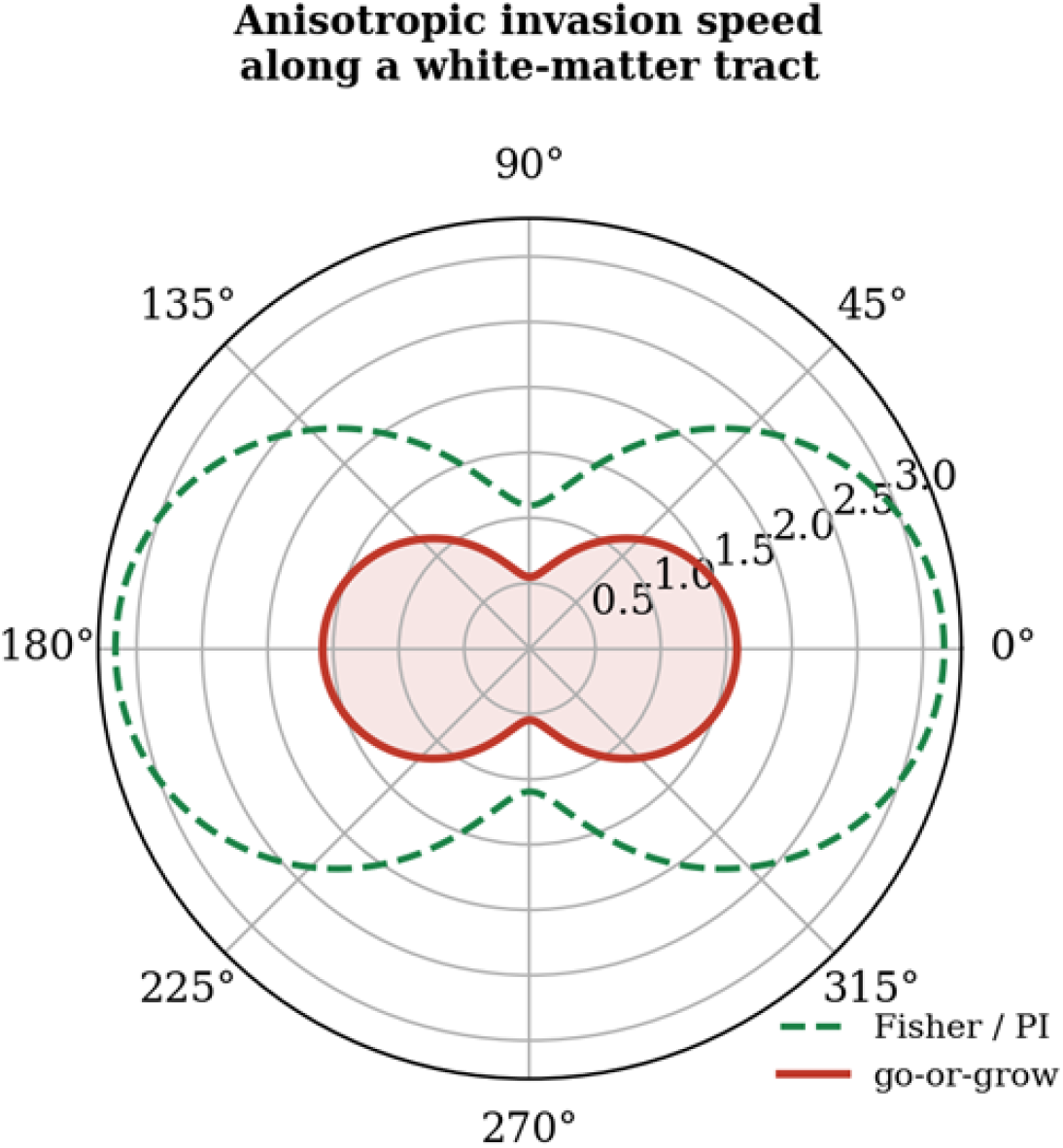
Anisotropy of invasion speed along a white-matter tract (polar plot; *D*_*∥*_ = 2.5, *D*_*⊥*_ = 0.3). The go-or-grow directional speed (Equation 15) (red) is a uniformly scaled-down copy of the Fisher/PI envelope (green) elongated along the direction of the fibre. Half-speed bound in all directions.

The implications of this mathematical discovery are enormous for patient-specific clinical modelling. Neurologists today have a specialized kind of MRI technique, called Diffusion Tensor Imaging, or DTI, which enables them to map the distinct white-matter “highways” in a given patient’s brain. Since the GoG penalty is proportional to the directional diffusion, the researchers can use raw images of the DTI scan from a patient and use those original directional tensors, **D**, directly in the GoG mathematical model, and simulate the precise, irregular 3D shape that the tumor will assume. Using the half-speed bound on these DTI models, clinicians would be able to predict these invisible subclinical invasion margins much more precisely, which could help to direct safer and more accurate surgical resections or radiation targeting.

## 4 Discussions

The effects of our work on the wider biological and clinical context are discussed in this section. We investigate the interplay between the phenotype balance and spatial constraint that not only determines the velocity of the invasive front, but also regulates the accumulation of the overall tumor mass at the macroscopic level. Additionally, we place these findings in the current paradigm of clinical imaging and discuss the potential merits and drawbacks of using the effective model based on a single parameter and the possible implications of reading out the internal phenotype balance as a guide for new, targeted therapeutic approaches.

The macroscopic invasion of glioma, determined by the GoG dichotomy of the individual cell, can be understood by means of an organizing principle: in the biologically relevant fast-switching regime, the invasion speed is bounded by half the classical Fisher speed, 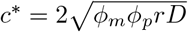.

However, a major clinical pitfall when using the traditional PI model to deduce intrinsic cell motility, *D*, from macroscopic imaging is the early detection of tumor invasion. The answer sharpens a bound recently proved for this model class by [12] [20, Theorem 9]. The migration stops temporarily during mitosis in order to allow the cell to divide, and thus the effective expansion speed of the tumor is intrinsically limited by the actual migration speed of the cells, meaning that the PI model inherently underestimates actual cell migration by mixing it into the migration fraction, *ϕ*_*m*_. The effects of the phenotypic transition rates (*α* and *β*) are highly visible in Fig. 8 (a). This figure depicts the final mass of a tumor with the same invasion speed as a function of phenotype balance in a 3D mass plot. In addition, Fig. 8 (b) shows that there is a near-perfect positive correlation between simulated front speed and final tumor mass (*r* = 0.98). As seen in this scatterplot, highly invasive tumors rapidly invade unoccupied tissue and are constantly releasing spatial constraints, thus allowing for a much larger overall tumor burden.

**Fig. 8.**
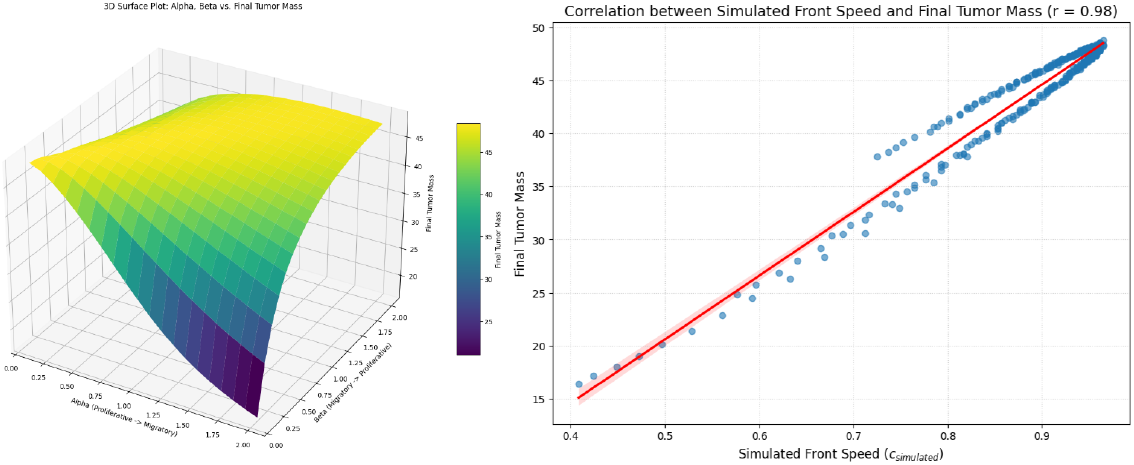
(a) Final tumour mass as a function of phenotypic switching rates in a topographical map. The surface shows the final tumour mass (Z-axis) for different values of the transition rates *α* (from Proliferative to Migratory) and *β* (from Migratory to Proliferative). The “migratory penalty” becomes evident by the steep gradient towards the low-mass valley at high *α* and low *β*, and the peak mass is located on the high-mass yellow plateau with proliferative bias; (b) Correlation between model invasion speed and final tumor size. The simulated front speed (*c*_simulated_) and the total final tumor mass are plotted against each other in a scatter plot, showing that there is a near-linear positive correlation (*r* = 0.98). The red line shows the linear regression fit with the confidence interval, with the red line indicating that maximising invasive potential also maximises the ability of the tumor to accumulate total biomass.

There is the existing literature on GoG models [20–24] with the authors Gerlee and Nelander deriving wave speeds for a related switching model [21, 23] and [22, 24] analysing traveling waves for go-or-grow glioma systems. The rapid switching down to the effective Fisher–KPP equation (Fisher-KPP) in Sec. 3.2 is a classical one [25]. It was recently shown by [12] that 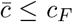 holds for a wide range of go-or-grow models, including the constant-rate model used here, by inserting one decay rate into the leading-edge dispersion relation. That argument gives an inequality bound but leaves two points to be clarified: that the bound is realized and that, if it is realized, what is the condition on the switching rates. We solve the dispersion relation exactly and find that the true minimiser, and the condition when it is attained, is the condition *ϕ*_*m*_ = , which is tight only at balance of phenotypes. On top of this sharpened result, we add: (i) the dimensionless discriminator 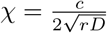, which translates the bound into a falsifiable test which can be computed from three quantities independently measurable, which, uniquely, reads the phenotype balance directly off the size of the shortfall observed; and (ii) the anisotropic, DTI-tensor generalization (sec. 3.4), which shows that the bound is geometry-independent and thus carries over unchanged to patient-specific, tensor-valued diffusion models, which the earlier traveling-wave literature for this model class did not consider.

[12] pose the question: what are the assumptions for the convergence of the fast-transition-rate system to its limit as *ϵ→* 0 in the case of traveling waves and what is the wave speed converging to when it does? We don’t solve it in the generality in which it is posed, that is, for general, possibly density-dependent transition functions *α*(*u, v*) and *β*(*u, v*), but there is direct numerical evidence for the case of constant rate, given in Fig. 6, which shows that the exact dispersion-relation speed in Sec. 3.1 approaches the closed form (Equation 10) as total switching rate increases, for every phenotype balance tested, with the half-speed bound being valid during the approach to the limit. This is provided as a specific instance of their general question; this is not the general resolution of their question. (*For more detail, please refer to the supplementary material section S3*).

## 5 Conclusions

Here, we will discuss our contribution by reducing many dense mathematical proofs and computer simulations to a handful of concrete biological truths. It provides the answer to the basic question that was asked at the start of the manuscript: **what does the microscopic behaviour of an individual cancer cell have to say about the macroscopic spread of a deadly tumor?**.

This half-speed limit is not merely some theoretical idea; it is a structural consequence, supported by multiple layers of rigorous proof:

- It was analytically derived by computing the so-called “Fisher-KPP limit” under the assumption that the switching between the states of the cells is very fast.
- The absolute minimum speed at which a mathematical wave can travel, known as the leading-edge dispersion relation, was used to verify the linear stability. We proved this by direct computer simulation (the heat maps and scatter plots you created).
- Importantly, they note, this speed limit is a consistent limit across the whole cell, regardless of how complex and anisotropic the geometry of the environment is, as the tumor travels through the brain’s white-matter tracts.

This is where the utility of this model comes into play: as a “read-out” of the hidden phenotypic state of a tumor that can be diagnosed. The half-speed bound 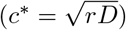 turns out to be a universal structural implication of the GoG dichotomy, but the power of this work is that it serves as a falsifiable scientific signature. We are successful in integrating front speed (*c*), cell motility (*D*) and proliferation rates in a unified model and offer a direct test of the GoG hypothesis to clinicians and experimentalists in real biological samples. However, if measured data of patient tumor properties are significantly off this predicted scaling relation, our hypothesis is thereby falsified, and we have a clear empirical route to further understand the behavior of gliomas. This model, therefore, represents a revolution in the understanding of an intractable phenomenon, switching from the phenotype to a signature that can be tested and whose parameters should not be confused with those of the phenotype. Finally, our simulations provide a data point toward an open question concerning fast convergence to wave speed raised by [12].

