## Supplementary for "Testable Clinical Signatures for Go-or-Grow Dichotomy of Gliomas along Anisotropic White-matter Tracts"

#### S1 Methods

##### S1.1 Governing Two-Phenotype Reaction–Diffusion PDEs

The model describes two types of mutually exclusive tumour cells, that is, one type that can migrate (migratory density $m(x,t)$) but do not proliferate, and another that can proliferate (proliferative density $p(x,t)$) but do not migrate. The sum of the local cell density is called the total local cell density, denoted by $\rho = m + p$. The following are the coupled equations:

$$\frac{\partial m}{\partial t} = D\frac{\partial^2 m}{\partial x^2} + \omega_{pm}(\rho)p - \omega_{mp}(\rho)m \tag{S1}$$

$$\frac{\partial p}{\partial t} = rp\left(1 - \frac{\rho}{K}\right) + \omega_{mp}(\rho)m - \omega_{pm}(\rho)p \tag{S2}$$

In this Equations S1 and S2, $D$ is the Fickian diffusion coefficient of the migratory cells, $r$ is the logistic growth rate of the proliferative cells, and $K$ is the maximum carrying capacity of the tissue. Switching rates between states are density-dependent. As cells become more crowded, the rate of proliferation to migration, $\omega_{pm}(\rho)$, increases, which pushes the cells to "go" and as cells have more free space, the rate of migration to proliferation, $\omega_{mp}(\rho)$, increases, which pushes the cells to "grow".

In Case 1, it is considered that $\omega_{pm}(\rho) = \alpha$, $\omega_{mp}(\rho) = \beta$ ($\alpha$ and $\beta$ are positive constants). It assumes that cells move between these two teams randomly at fixed linear rates and totally neglects local crowding. It is very amenable to analysis and has analytical information (baselines) that are clean. Case 2 reflects the true biological phenomena using non-linear Hill functions: $\omega_{pm}(\rho) = \alpha\frac{\rho^n}{\rho_c^n+\rho^n}$, an activation curve; switches on sharply once density clears a critical threshold $\rho_c$, and $\omega_{mp}(\rho) = \beta\frac{\rho_c^n}{\rho_c^n+\rho^n}$, an inhibition curve; shuts-off sharply as density climbs past $\rho_c$. Here, $n$ dictates the sensitivity (steepness) of the switch, and $\rho_c$ represents the threshold density

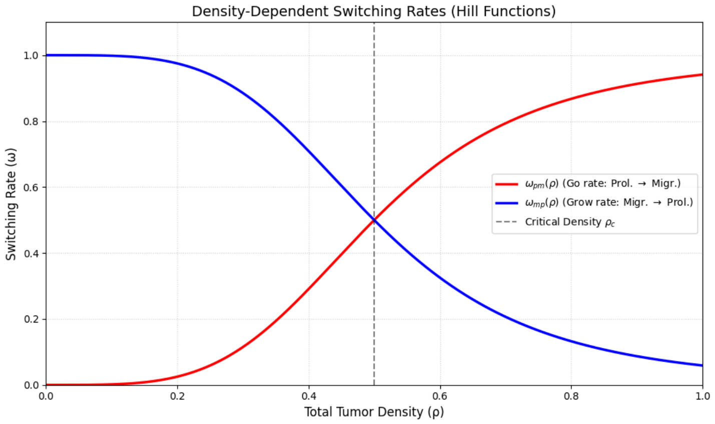


Figure S1: Phenotypic switching rates that are phenotypically density-dependent as a function of total tumor density $\rho$.

where cells become stressed. The escaping behavior due to crowding is modeled by the proliferative-to-migratory rate $\omega_{pm}(\rho)$ (red curve in Figure S1). In contrast, the migratory-to-proliferative rate $\omega_{mp}(\rho)$ (blue curve in Figure S1) corresponds to colonizing behaviour through free space that is inhibited by a Hill function. The point where these curves intersect is the critical density threshold $\rho_c = 0.5$, which defines the transition point at which a cell's dominant phenotype shifts from proliferation to migration. The steeper the slope of these curves, the greater the value of $n$; this is known as the Hill Coefficient ($n$). The higher the number of $n$, the closer the "Go-or-Grow" (GoG) decision will be to a binary switch; the lower the number of $n$, the closer the GoG decision will be to a gradual transition.

#### S1.2 Non-dimensionalisation of the Governing Equations

To facilitate numerical simulation and simplify the analytical derivations found in the main article, the governing two-phenotype reaction–diffusion equations are transformed into a dimensionless system. In this process the number of free parameters is reduced by scaling the variables by intrinsic biological scales of the tumor.

From the proliferation–invasion dynamics we define the following dimensionless variables for time $\bar{t}$, space $\bar{x}$, and the cell densities $\bar{m}$, $\bar{p}$, and $\bar{\rho}$:

- Time: Scaled by $1/r$ (proliferation rate), so that $\bar{t} = rt$.
- Scaled with the intrinsic invasion length scale, $\sqrt{D/r}$, so that $\bar{x} = x\sqrt{r/D}$.
- Density: Scaled by the maximum tissue carrying capacity $K$; $\bar{m} = m/K$; $\bar{p} = p/K$; and the density of the tissue is the total density

$\bar{\rho} = \rho/K$.

Applying these transformations, using the chain rule to the original dimensional PDEs, gives:

$$Kr\frac{\partial \bar{m}}{\partial \bar{t}} = D\left(K\frac{r}{D}\right)\frac{\partial^2 \bar{m}}{\partial \bar{x}^2} + \omega_{pm}K\bar{p} - \omega_{mp}K\bar{m}$$

$$Kr\frac{\partial \bar{p}}{\partial \bar{t}} = rK\bar{p}(1-\bar{\rho}) + \omega_{mp}K\bar{m} - \omega_{pm}K\bar{p}$$

By dividing both equations by $Kr$, the following fully non-dimensionalised system arises:

$$\frac{\partial \bar{m}}{\partial \bar{t}} = \frac{\partial^2 \bar{m}}{\partial \bar{x}^2} + \bar{\omega}_{pm}\bar{p} - \bar{\omega}_{mp}\bar{m}$$

$$\frac{\partial \bar{p}}{\partial \bar{t}} = \bar{p}(1-\bar{\rho}) + \bar{\omega}_{mp}\bar{m} - \bar{\omega}_{pm}\bar{p}$$

Through this scaling, the diffusion coefficient is scaled to $D = 1$; the proliferation rate to $r = 1$; and the carrying capacity to $K = 1$. As a result, the value of the reference classical Fisher speed ($c_F = 2\sqrt{rD}$) is naturally dimensionless. The dimensionless units are used throughout all of the simulated travelling-wave profiles and dispersion relation figures so that the physical geometry of the model can be universally applied.

#### S1.3 Mathematical Well-Posedness and Cooperative Comparison Principles

The physical formulation of the transition rates guarantees a mathematical structure of the system that is cooperative (quasi-monotone). The off-diagonal terms of the PDEs are strictly positive: the growth rate of the migratory cells, $m$, can only increase if the density of the proliferative cells, $p$, is increased, and vice versa. This cooperativity allows the use of standard parabolic theory to set up an invariant region. If the initial data $\rho(x, 0)$ is non-negative and satisfies the condition $\rho(x, 0) \leq K$ for any non-negative $x$, then the solution will be globally bounded and non-negative, that is, $0 \leq \rho(x, t) \leq K$ for all $t > 0$.

This is a property that involves an argument that is invariant, the rectangle, and the flux $\partial m/\partial t$ takes the negative sign at the strict boundary $m = 0$, the same as it does at the strict boundary $p = 0$.

Moreover, the maximum boundary line $\rho = K$ is a repelling barrier: the logistic proliferation term is strictly zero on it, whereas the terms accounting for the switching of phenotypes are strictly zero, conserving total density $\rho$. This formal comparison principle ensures that initial data that are ordered will be ordered over time, and that the speed at which the wavefront is pulled is well-defined according to the leading edge linearisation.

#### S1.4 Linearization

Linearization is a mathematical method that is applied to approximate the behavior of a complex non-linear system of ordinary differential equations (ODEs) around a particular point of interest, usually an equilibrium or steady state. This process starts in computational biology when the system's equilibria are determined by solving all the differential equations to find out where the biological populations ceases to change. After these steady states are found, the linearization determines exactly what the system will do in response to tiny "perturbations," like inoculating a small amount of tumor cells.

The tumor-free equilibrium ($m = 0, p = 0$) is evaluated using linearization to prove mathematically whether a small group of invading cells will survive or fail to colonize healthy brain tissue. The governing kinetic equations (for the local steady state analysis, neglecting spatial diffusion) are:

$$\frac{dm}{dt} = \alpha p - \beta m$$

$$\frac{dp}{dt} = rp\left(1 - \frac{\rho}{K}\right) + \beta m - \alpha p$$

We now take the partial derivatives of these equations with respect to $m$ and $p$ evaluated at $(0, 0)$ to create the Jacobian at the tumor-free state. Here, the total cell count is close to zero ($\rho \to 0$), the non-linear logistic term with the crowding parameter (1 - $\rho$/K) can be approximated to be equal to 1. The resulting Jacobian matrix $J$ is:

$$J = \begin{bmatrix} -\beta & \alpha \\ \beta & r - \alpha \end{bmatrix}$$

This tumor-free state is assumed to be stable to determine whether it is, we solve the characteristic equation $\det(J - \lambda I) = 0$ for eigenvalues ($\lambda$):

$$(-\beta - \lambda)(r - \alpha - \lambda) - \alpha\beta = 0$$

If the polynomial is expanded, it will result in a regular quadratic equation: The roots (eigenvalues) of this polynomial determine the stability of the tumor-free state. The transition rate ($\beta$) and the proliferation rate ($r$) are always positive values of the biology and thus the quadratic term has the constant sign of $-r\beta$. The properties of quadratics ensure that if $r > 0$ then the two eigenvalues have opposite signs, so the real part of one is strictly negative. The state without the tumor is mathematically unstable since there is a positive eigenvalue. That confirms that the brain tissue is unable to suppress the cancer, meaning that any small initial inoculum of glioma cells will inevitably grow, and will start the invasion front that spreads with the minimal speed $c^*$ you worked out in your leading edge analysis.

### S2 Results

#### S2.1 The Fast-Switching Asymptotic Reduction

The minimal invasion speed is mathematically obtained at the limit of fast switching of the system. This reduction will convert the two-phenotype system to a single effective scalar equation. The reduction makes the assumption that the time scales of the transition rates between migratory and proliferative states $\alpha$ and $\beta$ are much shorter than the spatial diffusion time scale and the time scale of cellular proliferation (Figure S2). For this limit, the local fractions of migratory cells ($\phi_m$) and proliferative cells ($\phi_p$) quickly reach a quasi-steady state characterized by the switching rates.

By applying this classical asymptotic approach to the governing PDEs, an effective Fisher–KPP equation, with an effective diffusivity $\phi_m D$ and an effective proliferation rate $\phi_p r$ is obtained. Therefore, the macroscopic invasion velocity is calculated in a closed form to be $c^* = 2\sqrt{\phi_m \phi_p r D}$, for this reduced system.

By maximizing the product $\phi_m \phi_p$ subject to the constraint $\phi_m + \phi_p = 1$, the exact mathematical minimiser is found. This shows that the maximum speed of the cells is $c^* = \frac{1}{2} c_F$ when and only when the time spent in both phenotypes is the same, i.e., $\phi_m = 1/2$. The fastest a GoG tumor can grow is half the classical Fisher speed, $\frac{1}{2} c_F = \sqrt{rD}$, which is the fastest a finite difference solution can grow (Figure S3).

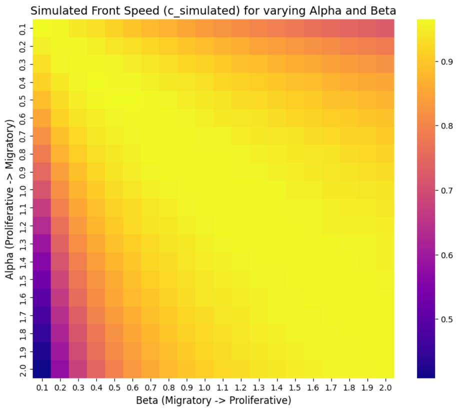


Figure S2: Simulated Front Speed $c_{\text{simulated}}$ for varying $\alpha$ and $\beta$

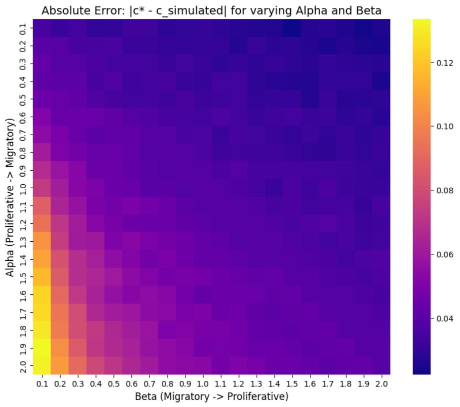


Figure S3: Absolute Error: $|c^* - c_{\text{simulated}}|$ for varying $\alpha$ and $\beta$

This is the "organising principle" of the tumor. It turns out to be a basic biological law - a tumor cell cannot move and divide together. Since the cell has to stop its motility machinery to divide, and then divide to move again, invasion depends heavily on the cooperation of different parts of the machinery. A cell that is 80% of its time moving covers a considerable distance, but no backup cells are born to advance the invasion front. On the other hand, if 80% of the time is spent on division, a huge army of cells is raised, but no one takes on new land. The maximum possible speed the tumor can have is when the cells spend 50% moving and 50% dividing.

##### S2.2 Density-Dependent Switching and Pushed Front Dynamics

The constant-rate formulation ($\omega_{pm} \equiv \alpha$ and $\omega_{mp} \equiv \beta$) produces a pulled front, which is determined by the properties of the linearisation of the front, at the leading edge of the front, but also allows density-dependent switching to be included, to capture complex microenvironmental behaviors.

The switching rates are defined as Hill functions: $\omega_{pm}(\rho) = \alpha\frac{\rho^n}{\rho_c^n+\rho^n}$, $\omega_{mp}(\rho) = \beta\frac{\rho_c^n}{\rho_c^n+\rho^n}$, where $\alpha$ and $\beta$ are positive constants. The "go" transition is an active process in these non-linear conditions and the growth of invaders is continuously provided from behind the leading edge. This internal density dependence fundamentally changes the invasion mechanics, causing the invasion front to more rapidly advance than it should, based on the leading edge of the linear model, and producing a "pushed" front. The pushed fronts are structurally sound in contrast to pulled fronts, which are very sensitive to stochastic variations and low cell-number cut-offs at the invasion boundary. This distinction in mathematics between pulled and pushed waves has clinical implications as to whether the tumor margin could be susceptible to targeted therapeutics that target the sparse subclinical cell populations.

##### S2.3 Derivation of the Anisotropic DTI Extension

Glioma invasion happens in a medium that is not homogeneous, but rather highly structured white-matter tracts of the brain, so the mathematics will need to be expanded to include spatial anisotropy. This enables the model to be calibrated directly from patient-specific DTI data. For this geometry, the value of the scalar diffusion coefficient is replaced by a tensor $\mathbf{D}(x)$ which is spatially dependent and reflects the local organization of white-matter fibres.

The migratory PDE is then transformed to become the equation,

$$\frac{\partial m}{\partial t} = \nabla \cdot (\mathbf{D}\nabla m) + \text{switching}$$

The system performs the state-of-the-art analysis for a planar invasion front propagating along a normal direction $\hat{n}$, which results in a direction dependent speed of the invasion front: $c^*(\hat{n}) = 2\sqrt{\phi_m\phi_p r\hat{n} \cdot \mathbf{D}\hat{n}}$. The same mathematical scaling factor, $\sqrt{\phi_m\phi_p} \leq 1/2$ works for all directions, and the directional Fisher speed is $2\sqrt{r\hat{n} \cdot \mathbf{D}\hat{n}}$, where $r$ is the number of points. It

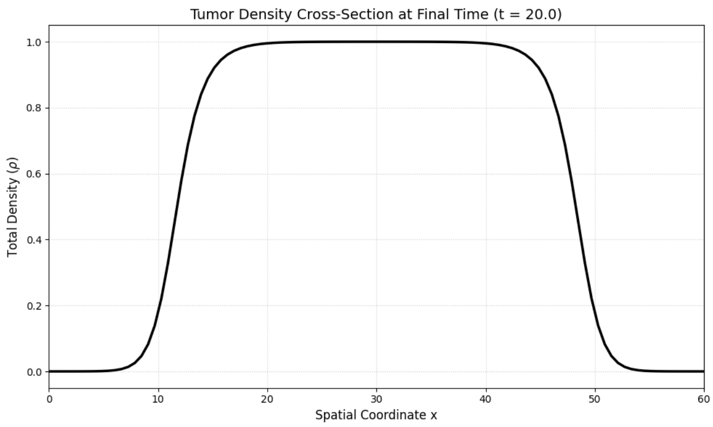


Figure S4: Macroscopic Traveling Wave Profile The 1D cross section of the total tumor density $\rho$ at the simulation time $t = 20.0$. The profile illustrates the case of a completely saturated tumour core, $\rho = 1.0$, growing with steep, symmetrical invasion fronts travelling at the theoretically predicted speed limit $c^*$.

demonstrates that the go-or-grow invasion envelope is a uniformly scaled down version of the classical Fisher envelope, which proves that the phenotypic speed bound is entirely geometry independent in complex tissue architectures.

### S3 Discussions

Diffuse gliomas grow invasively into normal brain tissue and surgery is never complete, making it virtually certain that the tumor will return. On the micro level, glioma invasion is controlled by a basic GoG: phenomenon—a phenotypic trade-off between distinct migratory and proliferative states of individual cells. In fact, biological evidence suggests that local microenvironmental factors, such as cellular crowding and free space, dynamically control the transitions between a dense core and sparse motile invasive edge, a behavior captured by a single-phenotype Fisher–KPP model of diffusion and proliferation. The scaling up of the single cell mutual exclusivity to determine the macroscopic invasion speed of the tumor margin is a key challenge in computational neuro-oncology.

The macroscopic invasion of glioma, determined by the GoG dichotomy of the individual cell, can be understood by means of an organizing principle: in the biologically relevant fast-switching regime, the invasion speed is bounded by half the classical Fisher speed, $c^* = 2\sqrt{\phi_m \phi_p r D}$, which is captured visually in Figure S5, which shows a spatial heatmap of the tumour's evolution from a localized seed to a large, radially expanding sphere in 20

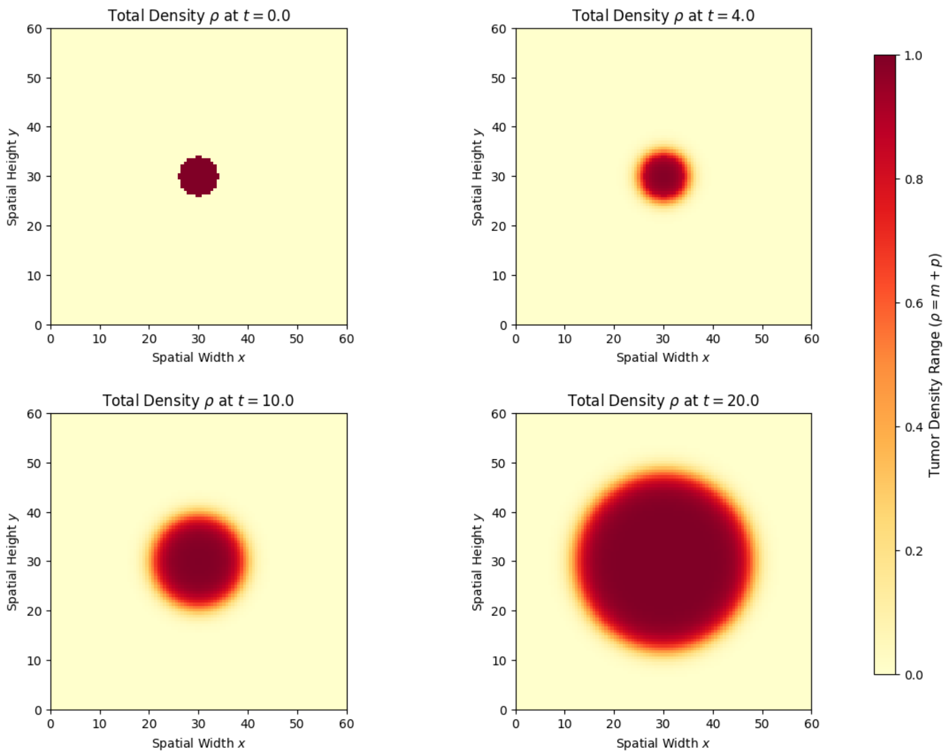


Figure S5: This 4-panel figure describes the macroscopic, radial expansion of a simulated glioma tumor over 20 time steps ($t = 0.0, 4.0, 10.0,$ and $20.0$). The total tumor density $\rho$ is depicted by a color gradient in a 60x60 spatial grid (Spatial Width $x$ vs. Spatial Height $y$). The light yellow color represents healthy tissue (density of 0.0), and dark red color represents the maximum tumour carrying capacity of a tissue (density of 1.0). At $t = 0.0$, a tumor exists as a very compact seed, and it quickly grows outward. It is quite striking how, at $t = 20.0$, it is all grown up into a large ball with a solid core of dark red and a clear, smooth outer growth of yellow-to-orange (invasive). Theories of travelling waves are given a visual representation here, with migratory cells making successful inroads into new areas and the main body continuing to multiply as it follows.

time units. The consequence of this mutual exclusivity in the structure is further illustrated in the density cross-section at $t = 20.0$ (Figure S4), which shows a travelling-wave structure with steep advancing edges travelling outwards at the theoretical $c^*$ speed limit, and a dense saturated core at the carrying capacity limit, $\rho = 1.0$.

The dynamic balance of this phenotype is of therapeutic importance, since therapies aimed at a particular subpopulation may affect the system. The ratio of migratory to proliferating cells is not fixed, as seen in Figure S5, it drops rapidly to $t = 4.0$ and then increases as the cells of the tumor core become crowded, and so they must take the motile “go” state to be able to find space.
